# Quantifying and Modelling Transfer Learning in Mice Between Consecutive Training Stages of a Change Detection Task

**DOI:** 10.1101/2019.12.19.883470

**Authors:** Ildefons Magrans de Abril, Marina Garrett, Douglas R. Ollerenshaw, Peter A. Groblewski, Shawn Olsen, Stefan Mihalas

## Abstract

Animals are known to be able to rapidly transfer knowledge between tasks with similar structure. We trained a set of mice on a visual change detection task with multiple stages, starting with direct transitions between gratings, adding an intervening gray screen, and subsequently moving to multiple sets of natural images. We observe that, when progressing to new stages, the performance increases very fast. However, when transitioning to a task of higher complexity, the peak performance decreases. This setup facilitates for the first time an experimental platform to study the transfer learning phenomena in mice using visual stimuli. Based on these results and additional neuroscience insight, we propose a cognitive model to explain the quick adaptation observed in mice. It extends a deep Q learning agent with a multi-tiered architecture and the possibility of performing a representation remapping at every level of the hierarchy to prevent the downstream propagation of representation anomalies. This architecture provides the substrate of an adaptation algorithm based on ideas of optimal transport of probability distributions. It matches well key behavioral aspects observed in mice and the experimental constraint that the mice are initially trained using a single task variant before it transitions to the new training phase. The modelling process helped us to gain biological insights: first, the optimal transport mechanism of the representation remapping indicated that a possible reason for the reduced performance when mice move to a new more complex task could be due to limited representation resources which were optimized for the previous task. Second, the multi-tiered architecture was first an engineering constraint and later a biologic insight confirmed by the literature. A final insight came from the computations required to perform the representation remapping. These computations are interesting because they could help to confirm this transfer learning theory by looking for similar neural correlates during the adaptation process.

## 1 INTRODUCTION

Transfer learning (TL) is the ability to apply knowledge learned in one context to another, thus speeding up the learning process. Different animal species present different levels of transfer learning ability. For instance, rodents trained to identify a triangle drawn with a solid line are unable to identify as the same object another triangle drawn with dots, while chimpanzees and humans can (Haskell, 2000a). However, rodents are feasible TL study models using auditory discrimination experiments (Kurt and Ehret, 2010) and active to passive avoidance (Sprott and Stavnes, 1974).

A recurrent observation in these studies is that TL is a context specific ability (Hobbiss, 2017). Even in humans there are clear documented examples where we fail to transfer between contexts. Hobbiss discusses a few (Hobbiss, 2017) like the ability of remembering a sequence of digits which does not predict ability to remember a sequence of letters (Tricot and Sweller, 2014), or that mastering brain-training games only makes people better at brain-training games (Simons et al., 2016). McGovern et al. (2012) investigate transfer learning in humans performing visual discrimination tasks. They studied the degree of transfer between three related discrimination tasks where all three tasks require the participant to identify the orientation and/or the curvature of the elements. Results demonstrated transfer learning from one task to another provided provided they share a common cue for their processing (e.g. subjects trained on orientation discrimination showed improvement on curvature related tasks as a function of the relative complexity between train and test tasks). Similarly, in rodents performing auditory discrimination learning, transfer is observed to depend on task difficulty (Kurt and Ehret, 2010).

Although there is not an accepted comprehensive theory to explain transfer learning (Haskell, 2000b), recently a few authors have proposed cognitive models to explain related phenomena like task generalization and continuous learning of many tasks. Wang et Al. (Wang et al., 2018) discuss the interplay between the prefrontal cortex (PFC) and the dopaminergic reinforcement learning system (DS). In this model, the DS is responsible for the specialization of the PFC for a particular set of tasks. Then, the trained PFC model is able to operate as a specialized RL system able to transfer the acquired skills to new similar tasks. It is interesting that this model offers a theory to explain experimental evidence of both model-based and model-free RL in the brain and it explains the observation of dopamine like error signals in the PFC. On the other hand, this model requires a large number of similar tasks to train the specialized RL system. Thus, it does not explain the quick adaptation observed in animals to a similar task when they have been previously trained on very few related tasks (see section 2 for an example where transfer learning is observed after training the animal on a single task).

Yang et al. (2019) propose a relevant computational platform to study the ability to perform many tasks. They noted the duality in neuron function specialization and its consequences in the brain strategies to implement transfer learning. On one hand, it is known that functional specialization is a design principle of the brain. On the other hand, mixed selectivity neurons are commonplace in the PFC. They theorize that specialization is desirable when a particular computation is beneficial to several behaviors. In this case, transfer learning could be achieved through a compositional approach. On the other hand, mixed functionality is hypothesized to be encouraged by the need to maintain future flexibility. In this second case, the continuous process of refining the neural function would be substrate of the transfer learning ability.

Although the purpose of this paper is to propose a cognitive model not a machine learning method, we hope the ideas presented in this paper can inform future work in machine learning. Supplementary section 1 discusses a generalization of our model that we believe could also serve as an starting point to develop a new transfer learning approach. In that sense, we briefly review main approaches which partially share key design decisions with our model. Zhang et al. (2018) propose a reinforcement learning agent with two main architectural components: the environment dynamic module and the reward function. They show they can re-train them independently, thereby speeding up the learning process by transferring the still relevant module to the new task (Zhang et al., 2018). Another relevant machine learning approach is to use optimal transport for domain adaptation in the context of supervised classification problems (Courty et al., 2017).

This paper is organized in three main blocks. Section 2 quantifies the transfer of task related knowledge when mice are trained with a curriculum of increasingly complex change detection tasks using visual stimulus. Section 3 discusses a conceptual model and its implementation details to explain the quick adaptation observed in mice. This model is motivated by our own analysis in mice, as well as existing experimental data in mice and other animals. Finally, section 3.4 presents the model simulation results with respect to mice behavior.

## 2 ANALYSIS OF TRANSFER LEARNING IN MOUSE

We use data from the visual behavior dataset from the Allen Brain Observatory, standardized in vivo survey of physiological activity in the mouse visual cortex while performing a change detection task. We train each mouse following a sequence of task variants. Mice advance to more challenging training phases when they are able to reach a maximum *d*′ (metric of performance described below) larger than 2 for at least 2 sessions out of the last 3, where each session last for 1 hour and consists of an average of 400 trials. Before starting the regular training protocol, mice are exposed to 15 minutes of associative trials with the same trial format used in training stage 1 but with a guaranteed reward for every change. Figure 1 shows a diagram of the training protocol with four training phases.

**Figure 1.**
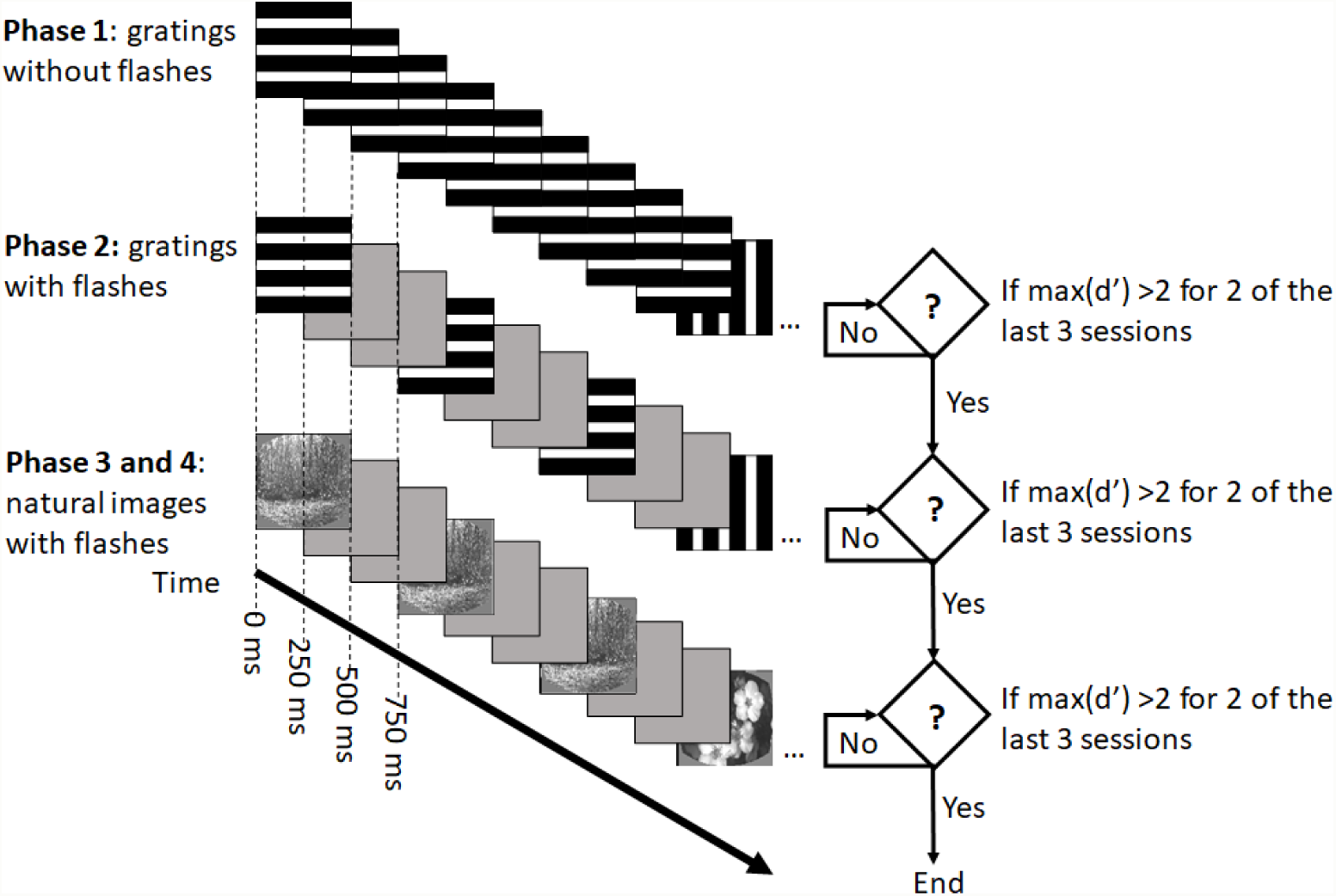
Training protocol. Square images represent full screen stimuli.

- Training stage 1: mice are presented a sequence of images with horizontal and vertical gratings at 100% contrast. These gratings are static so they are presented continuously with changes occurring according to a truncated exponential distribution of change times. Mice are rewarded when they respond following a transition between horizontal and vertical gratings.
- Training stage 2: mice are also presented a sequence of images with horizontal and vertical gratings. In this case, each image is presented for 250 ms followed by a gray scale screen for 500 ms.
- Training stage 3 and 4: the last two stages are similar to stage 2 but now mice are presented with natural images instead of gratings chosen from a dictionary of 8 images. Stages 3 and 4 use different image dictionaries.

Figure 2 contains additional task descriptions. There are two types of trials: *catch trials* have the same image repeatedly presented while *go trials* present a different image in the last step of the trial. Besides the trial types, figure 2 describes additional concepts: an *aborted trial* (A) happens when the mouse licks before the scheduled image change, a *hit* (H) is when a mouse licks in a 600 ms window following an image change in a go trial ^1^, a *false alarm* (FA) happens when the mouse licks during the same time window of a catch trial, a miss (M) occurs when the mouse does not lick at all during a go trial and correct reject (CR) happens when when the mouse does not lick at all during a catch trial. Given these definitions we define *d*′ as follows:

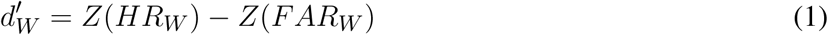

where Z is the is the inverse of the cumulative distribution function of the Gaussian distribution and *HR*_*W*_ and *FA*_*W*_ are the hit rate and the false alarm rate within a window of *W* non-aborted trials^2^, bounded to the interval 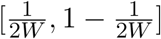.

**Figure 2.**
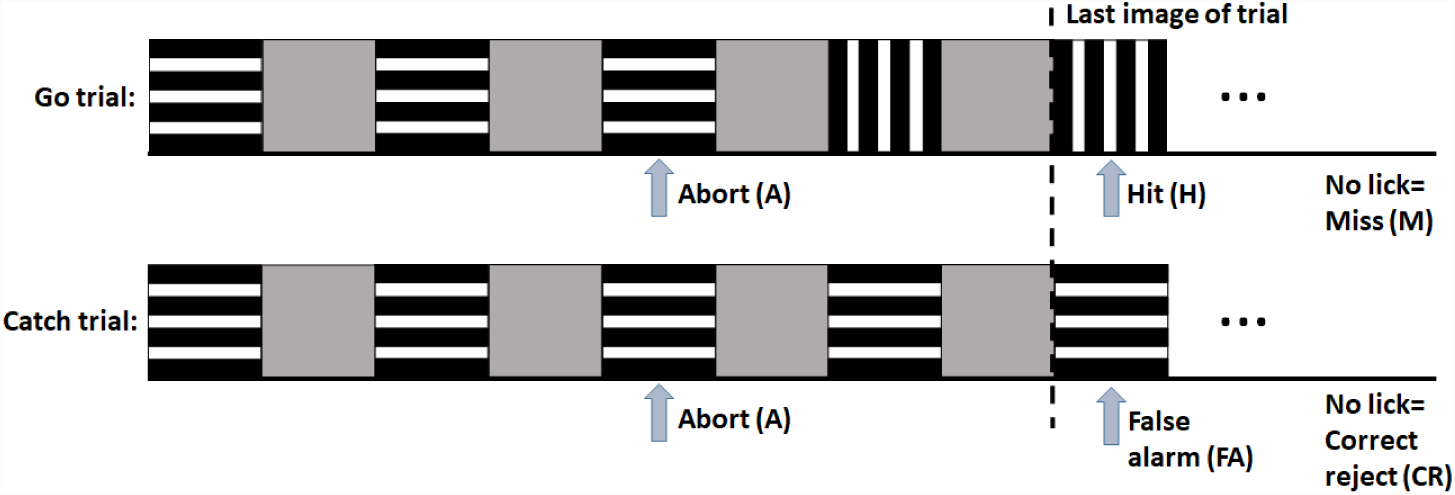
In the task, there are two types of trials: Go and Catch corresponding to trials with a change or without respectively. Depending of the mouse response, a trial can be classified as: i) Hit: when a mouse licks on the last image of a go trial, ii) False alarm happens when the mouse licks during the last image of a catch trial, iii) Miss occurs when the mouse does not lick at all during a Go trial and iv) Correct reject happens when when the mouse does not lick at all during a catch trial

### 2.1 Quantifying transfer learning

We study the transfer learning capacity of 27 mice at different stages of the training procedure discussed in section 2. We quantify the learning efficiency of the mice by first computing the time series of running *d*′, with window (W) of 100 trials, for each mouse and each session of every training phase. The second step is to identify, for each first session of every training phase *j*, the first trial number where *d*′ is equal or greater than 1 (*t*_*j*_). Finally, we quantify the mouse transfer learning ability by measuring the relative learning efficiency when it is exposed for the first time to training phases 2, 3 and 4 compared to the learning efficiency demonstrated during the first session of the first training phase. This rate is formally defined as follows:

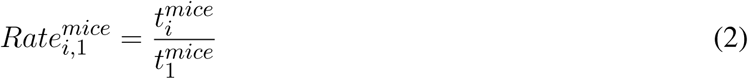

Table 1 shows the median and corresponding median absolute deviation of each 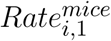 indicator. The main observation is that mice are consistently reaching a *d*′ greater or equal than 1 during the first session of training phases 2, 3 and 4 more quickly than during the first session of training phase 1. This indicates the mouse’s ability to quickly accommodate to task variants by leveraging skills acquired in previous tasks. The 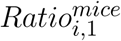 indicator is used later in section 3.4 to discuss the plausibility of a new transfer learning model and to compare it against a baseline implementation. Mice rates are significantly below 1 with p-values^3^ of 0.00049, 0.0004 and 0.006 respectively.

**Table 1.** Median and median absolute deviation of 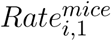 for i ∈ {2, 3, 4} and a population of 27 mice.

| Median (median absolute deviation) | $Rate_{2,1}^{mice}$ | $Rate_{3,1}^{mice}$ | $Rate_{4,1}^{mice}$ |
| --- | --- | --- | --- |
| | 0.10 ($\hat{\sigma} = 0.10$) | 0.22 ($\hat{\sigma} = 0.18$) | 0.17 ($\hat{\sigma} = 0.22$) |

Figure 3 shows the evolution of the average *d*′ during particular training sessions of interest to study transfer learning. Figure 3.a) corresponds to the first session of the first training phase. During this session, mice are exposed to the change detection task for the first time. We will use this case as reference case to quantify the mouse transfer learning ability. In this case, it takes an average of 638 trials 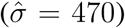 to achieve a *d*′ of 1. Figures 3.b), c) and d) illustrate the learning efficiency of the mouse during the first session in a new training phase just after graduating from the previous training phase. The first observation from figures 3.b), c) and d) is that mice are able to learn faster during the first session after transitioning to a new training phase than during the first session of the first training phase. Precisely, our mouse population is able to reach a *d*′ of 1 in just an average of 83 trials 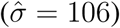 when transitioning from phase 1 to 2, an average of 114 trials 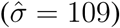 from phase 2 to 3, and an average of 133 trials 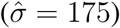 from phase 3 to 4.

**Figure 3.**
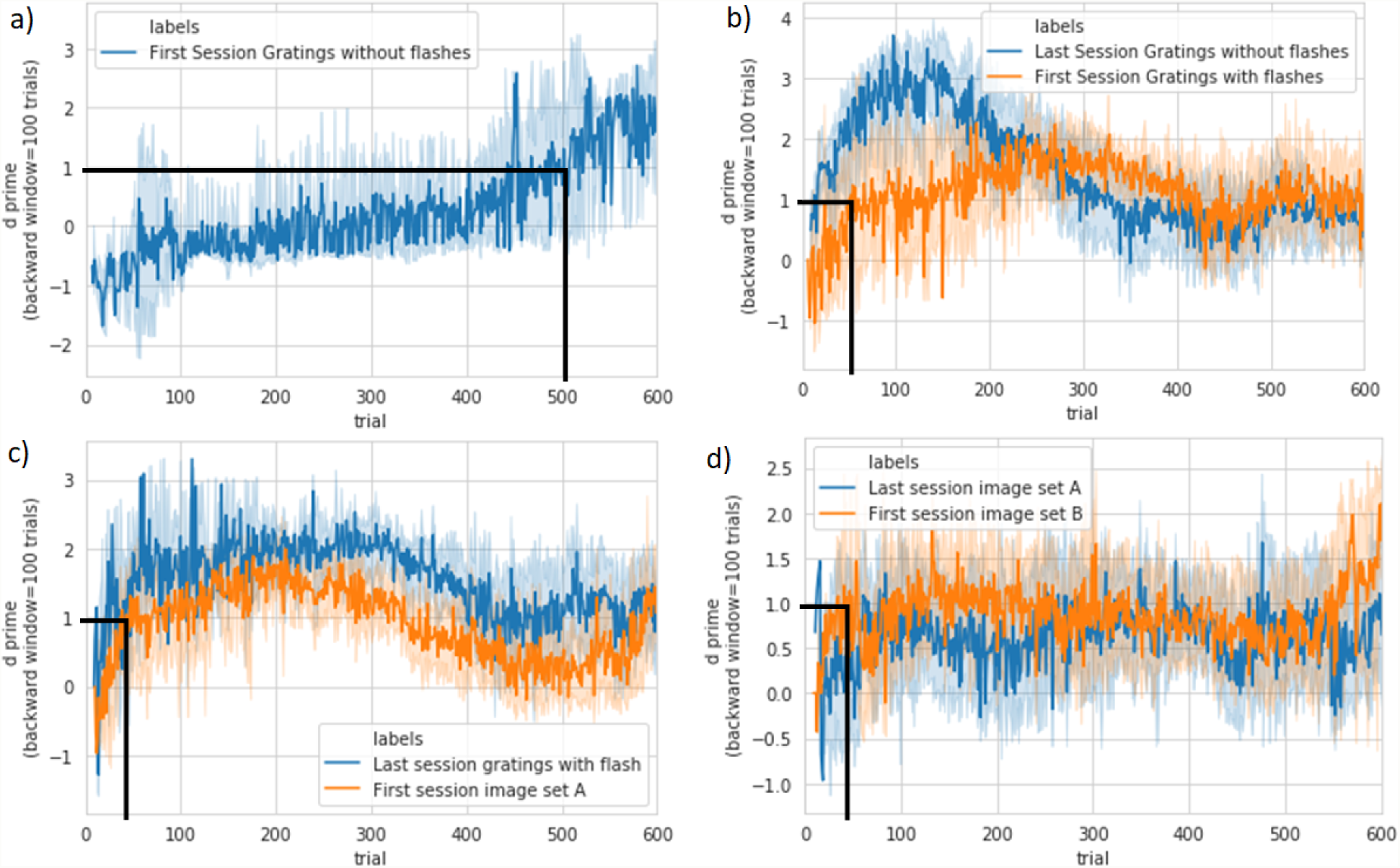
Average *d*′ for a population of 27 mice. (a) Reference case: first session of first training phase. (b) Blue: last session of first training phase, orange: first session of second training phase. (c) Blue: last session of second training phase, orange: first session of third training phase. (d) Blue: last session of third training phase, orange: first session of fourth training phase.

Figures 3.b) and c) show also that when mice move from phase 1 to 2 and from 2 to 3, they reach a maximum average *d*′ lower than during the last session of the previous training phase. These two training phase transitions correspond to a raise in representation complexity requirements: when moving from phase 1 to phase 2, we add grey images for 500 ms after showing gratings for 250 ms, thus adding complexity to the temporal dynamics of the task; when transitioning from phase 2 to phase 3, we are increasing the size of the image dictionary from 2 types of gratings to 8 different natural images. We hypothesize that this phenomena could be due to a quick adaptation mechanism trying to recycle existing resources to speed up the learning process. Therefore, the increased representation complexity in the new task after the transition could cause a slightly reduced performance. This hypothesis is confirmed by the analysis of the transition between phase 3 and 4. The difference between these two phases is that the image dictionary changes to a new one with the same number of images, thus the representation complexity requirement is the same. Figures 3.d) shows that the maximum average *d*′ during the first session of training phase 4 is not lower than during the last session of the previous training phase.

These results indicate that mice performing visual discrimination experiments are able to take advantage of knowledge acquired during their recent training history to improve their learning efficiency of new task variants. A second observation is that the maximum achievable performance due to the quick adaptation mechanism seems to depend of the relative representation complexity requirements of the new task.

## 3 MODEL

The model architecture is based on two architectural principles: i) a multi-tiered architecture and ii) the idea of distributed representation firewalls. Each principle is in turn motivated by relevant neuroscience evidence presented during the background discussion and new experimental results presented in section 2 which are later validated against the model simulations.

Our first design principle is a multi-tiered architecture. It is first suggested by the well-known hierarchical organization of the sensory cortex. A more precise indication of the relevant hierarchy is given by in-vivo recordings from monkeys trained to detect changes in patterns composed of vibrotactile flutter stimuli. These recordings show that the primary somatosensory (S1) has a faithful representation of stimuli but not enough to decide whether there is a change while representation in dorsal premotor cortex (DFC) codes specific combinations of past and present stimuli and are strongly predictive of the monkey’s behavior (Rossi-Pool et al., 2016).

The second design principle is based on the the hypothesis of the existence of representation firewalls in the input of each tier. They are responsible for the conversion of actual inputs to expected ones, thus preventing downstream encoding anomalies. Even though we move our eyes frequently, our perception of the world is stable. While the initial representations in the visual system are in retinotopic coordinates, these get transformed to body or world centered coordinates. Such transformations are similar in concept to the representation firewalls we propose here. Just as allowing such representation remapping following saccades helps with identifying an object, similar remapping following representational shifts which otherwise keep the structure of the world the same help in fast learning. An example of representation shifting phenomena is observed in monkeys trained to classify 360° of visual motion directions in two categories. After monkeys were retrained to group the same stimuli into two new categories, lateral intraparietal neurons shifted towards the new category membership, while neurons in middle temporal areas kept the same direction selective representation (Freedman and Assad, 2006). A second example is neuron receptive fields in macaque area V4 shift towards a new target even before initiating the saccadic movement towards this target (Tolias et al., 2001). We speculate that the hippocampus could also play an important role in the implementation of the representation firewall functionality. Evidence in that direction is two fold: i) both temporal and spatial coding can be observed differently in some hippocampal regions (review paper (Eichenbaum, 2017)) and ii) evidence of grid cells encoding statistics of 2D space (Dordek et al., 2016; Whittington et al., 2018). The computation of probability masses of new and original input spaces is a functional requirement of the actual adaptation algorithm that we will discuss in section 3.3. The multi-tiered architecture tailored to change detection tasks together with the idea of representation firewalls are our main model architectural biases. In the introduction, we discussed that transfer learning seems to be a context specific ability. Our particular multi-tier architecture also facilitates transfer learning across a finite range of task variants. Supplementary section 1 discusses a generalization of this model to accommodate more complex environments. Next, we will discuss the model inference and adaptation details, the simulated environment and the experimental results.

Figure 4 presents the model architecture. We assume a discrete action space (change/no change) and a continuous fully observable state space, where observations are the images presented to the mouse every time step *i*_*t*_. Input images are pre-processed by a pre-trained convolutional neural network (Targ et al., 2016). The CNN output is a 84 dimensional feature vector which acts as a model input state *s*_1,*t*_. The model output is a vector which assigns a quality value to each state-action pair *q*_*t*_. Our action selection is 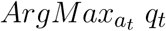 with an exploration noise *∈*.

**Figure 4.**
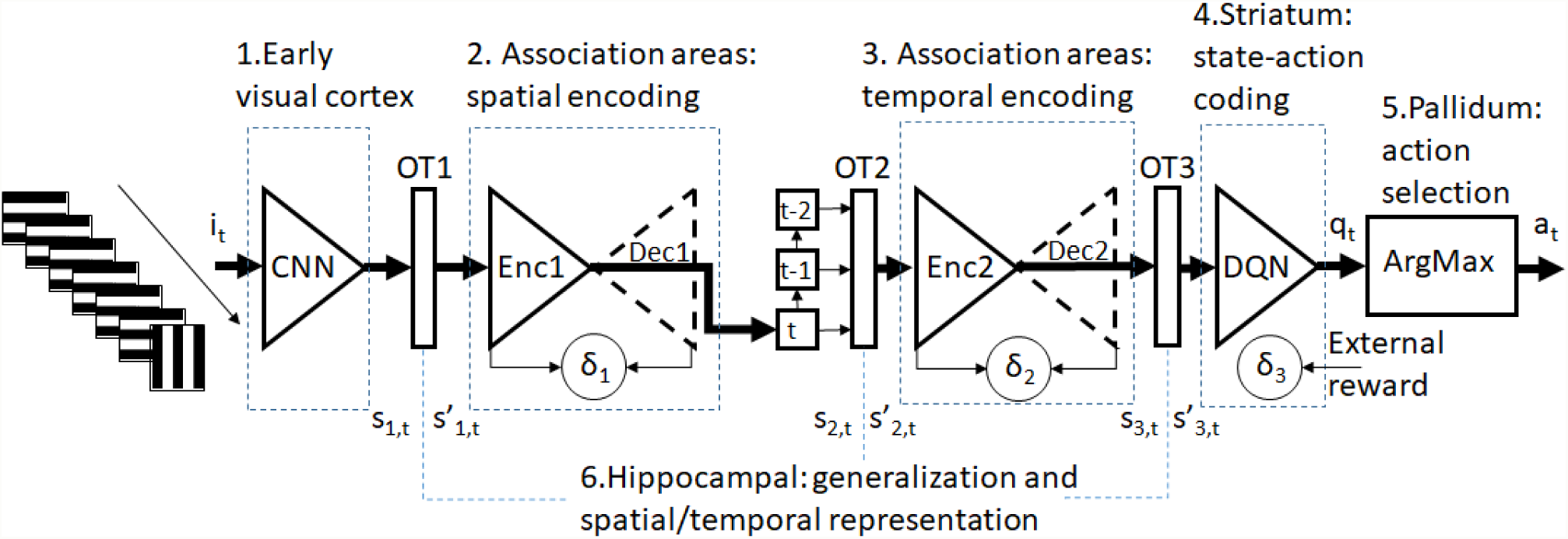
Model overview. Table 2 defines all the nomenclature of the figure.

Our model implementation has three tiers and each tier has a similar functional structure with 3 functionalities. The model tiers are *Enc*1, *Enc*2 and *DQN*. Each tier has a function *OT* which transforms the input and is responsible for domain adaptation 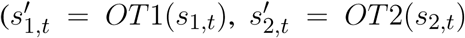 and 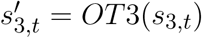. Figure 5 shows a conceptual diagram of this functionality. Section 3.3 discusses this key component.

**Table 2.** Nomenclature used in figure 4

|  |  |
| --- | --- |
| $i_t$ | Stimulus data |
| $s_{1,t}$ | Feature vector generated by the CNN (feature vector in tier 1) |
| $s'_{1,t}$ | Remapped feature vector in tier 1 |
| $s_{2,t}$ | Feature vector generated by the tier 1 encoder |
| $s'_{2,t}$ | Remapped feature vector in tier 2 |
| $s_{3,t}$ | Feature vector generated by the tier 2 encoder |
| $s'_{3,t}$ | Remapped feature vector in tier 3 |
| $q_t$ | Quality value for each possible action |
| $a_t$ | Selected action |
| $OT_{[1,2,3]}$ | Remapping function in level 1,2 and 3 respectively |
| $\delta_{[1,2]}$ | Reconstruction loss of autoencoder in level 1 and 2 respectively |
| $\delta_3$ | External reward |
| CNN | Convolutional neural network |
| Enc1 | feature extraction in tier 1 (also encoder of autoencoder of tier 1) |
| Dec1 | Decoder of autoencoder of tier 1 |
| Enc2 | feature extraction in tier 2 (also encoder of autoencoder of tier 2) |
| Dec2 | Decoder of autoencoder of tier 2 |
| DQN | FFN delivering a quality value for each possible action |
| ArgMax | Action selection |

**Figure 5.**
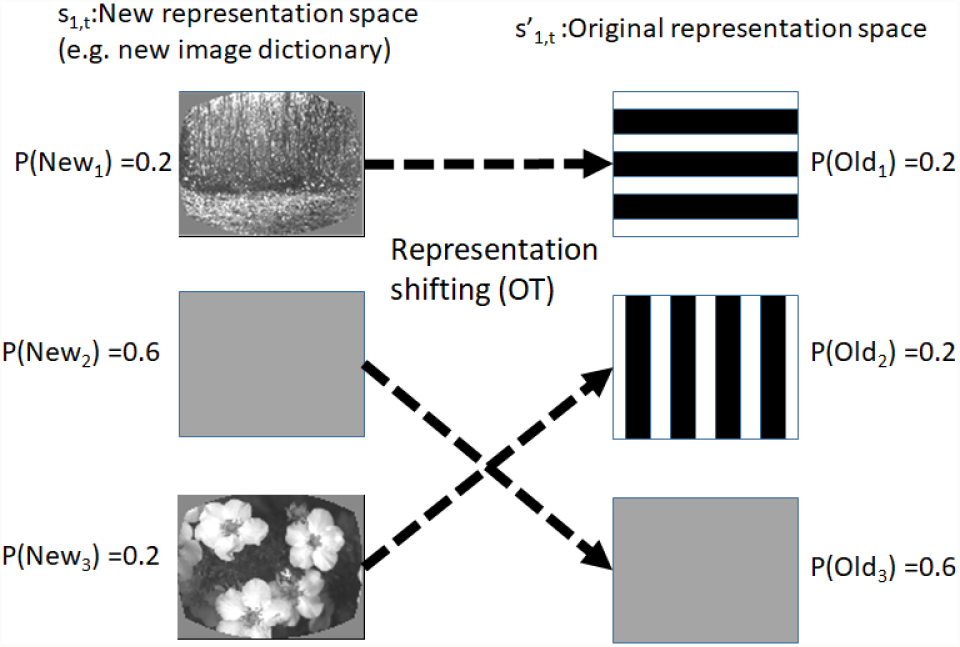
Concept of representation remapping (OT).

*Enc*1 does feature extraction from current image frame and reduces dimensionality from 84 to 6 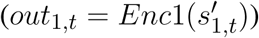. The second tier is *Enc*2. It extracts dynamic environment features. We are unsure of the exact location for these processes, and it could be in higher order visual or association areas. They are both implemented with Feed Forward Networks (FFN). *Enc*2 uses the current and the 2 previous *Enc*1 outputs as inputs 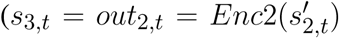 where *s*_2,*t*_ = *Concatenation*(*out*_1,*t*_, *out*_1,*t*−1_, *out*_1,*t*−2_)). Finally, the policy network *DQN* is also a FFN which computes the quality vector 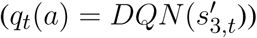. Finally, the agent selects the action (lick or not lick) with bigger quality ^4^.

Each tier has 3 functionalities:

- i) Feature extraction as discussed above.
- ii) Anomaly detection in the input distribution.
- iii) Representation remapping through optimal transport.

These three functionalities facilitate the quick adaptation to environment variants. Feature extraction is implemented using respective FFNs (i). The second functionality is to detect input anomalies. *Enc*1 and *Enc*2 implement this functionality by fitting a decoder *Dec*1 and *Dec*2 respectively and then using the reconstruction loss to monitor possible changes in the input distribution of the corresponding tier:

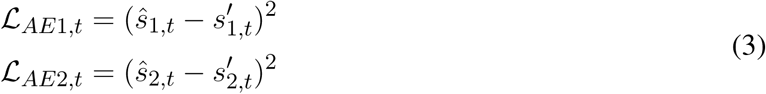

Anomalies in the policy network *DQN* are identified with a drop in agent running performance. A sustained anomaly in time is indication that the input distribution of a tier has changed. We updated these indicators continuously as follows:

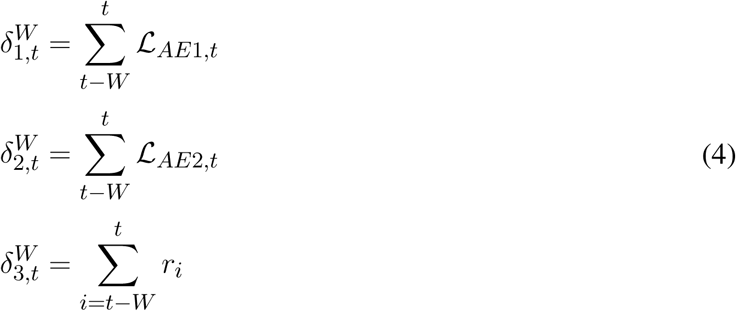

where W is a backward window and *r*_*t*_ is a reward delivered by the environment. We discuss the environment emulation details in section 3.1.

The third functionality is the representation remapping through optimal transport. When both source and target distribution are discrete, then the optimal transport is defined as:

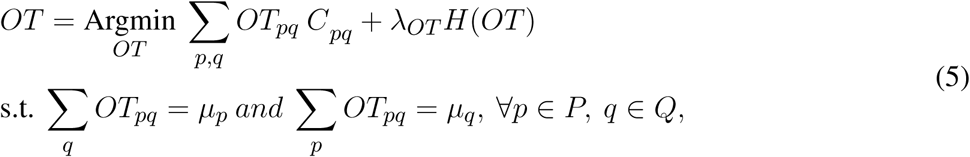

where *P* and *Q* are the categories of the new and the original input distributions, *OT* and *C* are the coupling matrix and the transport cost matrix respectively, *OT*_*p,q*_ indicates the amount of probability mass of category *p* in the new input distribution that is transported into category *q* in the original one, *µ*_*p*_ and *µ*_*q*_ are the probability masses of the categories *p* and *q* from the new and the original distributions respectively, *H*(*OT*) is the entropy of the transport matrix, and *λ*_*OT*_ controls the the smoothness of the transport matrix. By increasing *λ*_*OT*_ we can encourage a sparse *OT* which is interesting in case of a deterministic environment change (e.g. color shift) and by reducing *λ*_*OT*_ we can accommodate an stochastic environment change (e.g. variation of luminosity due to random clouds).

Equation 5 is a convex optimization problem that depends on the probability masses *µ*_*p*_ and *µ*_*q*_, and the transport cost matrix. This formulation has two main difficulties: first, it requires a online unsupervised process running in parallel to cluster the input space and to extract the probability masses as well as the cluster prototypes *P* and *Q* from the new and the original input data respectively. In this paper we assume that this process occurs but we will not discuss a particular solution (e.g. (Lin, 2013)). Second, the cost matrix does not exist. Instead, we propose to evaluate the cost of a particular transport by measuring the external reward. For instance, if we are trying to perform adaptation on *Enc*1 input, then we will aim to find an *OT*_1_ matrix that minimizes −*δ*_*DQN,t*_ while taking into account the probability mass preservation and overall entropy constraints. The following cost function implements our goal to transform the new input distribution into the old one while preserving the environment dynamics and the agent task performance:

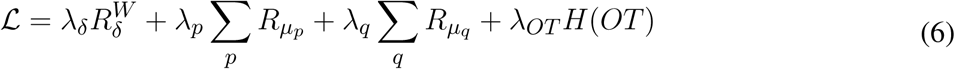

where:

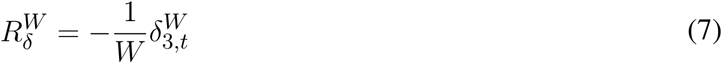

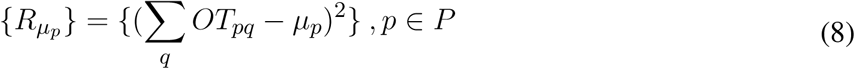

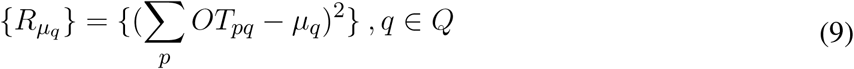

For the original distribution, probability mass estimates are computed using data captured during normal operation. For the new distribution, estimates are computed using data captured during the anomaly detection period. In future work we would like to extend equation 7 with reconstruction losses from the tier 1 and 2. This could speed up the learning because 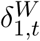 and 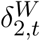 do not depend of sparse external rewards:

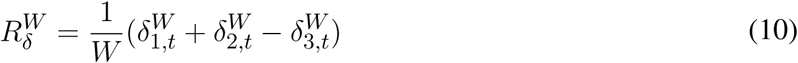

### 3.1 Environment emulation

Due to the continuous flow of the change detection trials, we emulate this environment as a single continuous process made of a sequence of trials and each trial in turn consists of a sequence of images. The exact format of the trial image sequence as a function of the training phase was discussed in section 2. The emulated environment issues a reward every step as follows:

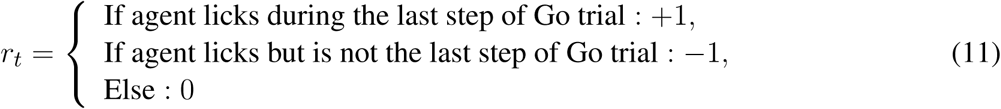

This reward function assumes that being able to drink water when trying is used by the mouse as a positive reinforcement and that not being able to do so when trying is a negative one. We are aware that the reward functions is to some extend arbitrary in the sense that we don’t fully know what is the internal reward signal driving the mouse learning process. To minimize this uncertainty, we have summarized the effect of different negative rewards in the agent performance (see supplementary section 2). We have compared the agent time required to reach d’ of 2 by training the DQN tier starting from random weights, using a dictionary of 3 different natural images and 4 different reward functions which uniquely differ in the negative reward −1, −0.2, −0.1 and 0. Results show that the negative reward does not have a well defined effect on the agent performance. Therefore, choosing a negative reward equal to −1 would not bias the results.

### 3.2 Model inference

Initial model inference of the three tiers is done sequentially keeping *OT* functions as identity functions. First, *Enc*1 is fit as part of a first autoencoder. Second, we train *Enc*2 as part of a second autoencoder after connecting the output of *Enc*1 to the input of *Enc*2. Autoencoders weights are optimized using the loss functions ℒ_*AE*1,*t*_ and ℒ_*AE*2,*t*_ respectively as defined in equation 4. Finally, we train the policy network *DQN* after connecting the output of *Enc*2 to the input of *DQN*. This initial model inference is done using trial sequences from the simplest training phase discussed in section 2 (i.e. gratings without grey flashes).

The training of *DQN* could be done with any reinforcement learning algorithm. The particular choice does not affect the quick adaptation functionality which is the main contribution of this paper. Rewards are granted according to the reward function discussed in section 3.1. We used a deep Q learning algorithm (Mnih et al., 2013) with discount factor *γ* = 0 (only the immediate reward is taken into account to update *DQN*), constant exploration noise, replay buffer with experience replay and Huber loss defined as:

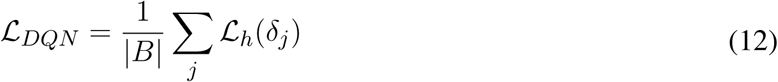

where:

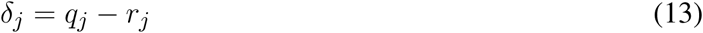

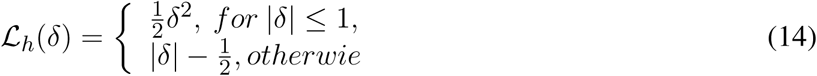

### 3.3 Model adaptation

The model adaptation phase starts when the architecture is well trained (see section 3.2), under operation and some of the indicators, presented in equation 4, detect an anomaly. The main result of this section is algorithm 2. This algorithm describes the representation remapping process in detail assuming that we know the tier where we want to do the intervention. We also discuss a candidate algorithm to identify the tier level but we have not used it in the simulations.

Algorithm 1 is responsible for deciding what intervention to perform as a function of the current state of the anomaly detection indicators. There are three possible interventions (*OT* 1, *OT* 2 and *OT* 3) corresponding to an optimal transport operation on the input distribution of *Enc*1, *Enc*2 and *DQN* respectively. The optimal transport operation is defined by a transport matrix with *P* rows and *Q* columns where *P* and *Q* are the number of clusters identified in the new and the original input distribution respectively.

#### Algorithm 1: Overall algorithm to detect and decide where to perform a representation remapping

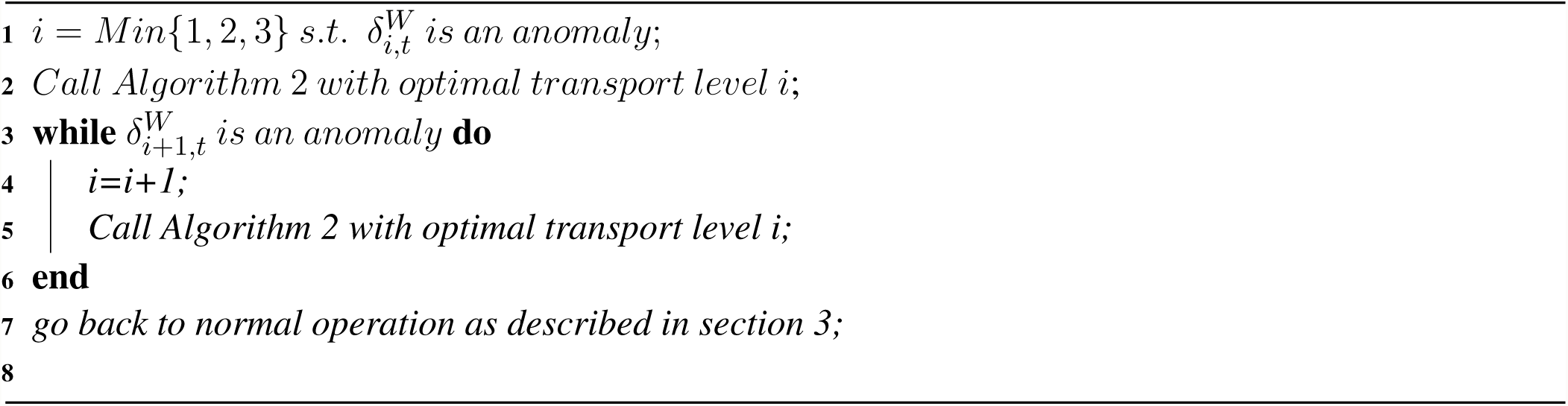

The intervention selection algorithm is based on the idea that an anomaly at a particular level will start an anomaly cascade in all downstream levels. Therefore, if we want to solve the cascade we have to start by trying to correct the top level anomaly. When the top-level intervention is not able to correct the anomaly cascade, the algorithm proposes a follow-up intervention one level below. This can happen for at least two reasons: 1) the new input distribution has a larger representation complexity than the original one and 2) the intervention at the top-level is not enough because there are indeed 2 simultaneous changes for instance a change in the image dictionary together with a change in the environment dynamics (e.g. adding an additional grey image every other image).

All three possible interventions proposed by algorithm 1 are executed by the same algorithm 2. A key difficulty to design algorithm 2 is that agent actions depend of agent observations and how those are transformed by *Enc*1, *Enc*2, *DQN* and the current state of the corresponding transport matrix. In particular, a bad transport matrix would make it very hard for the agent to collect environment rewards *r*_*t*_. To cope with this difficulty, we propose an online optimization algorithm which alternates the interaction with the environment in order to collect /observation/action/reward traces with the progressive optimization of the transport matrix.

#### Algorithm 2: Monte-Carlo based optimal transport for domain adaptation

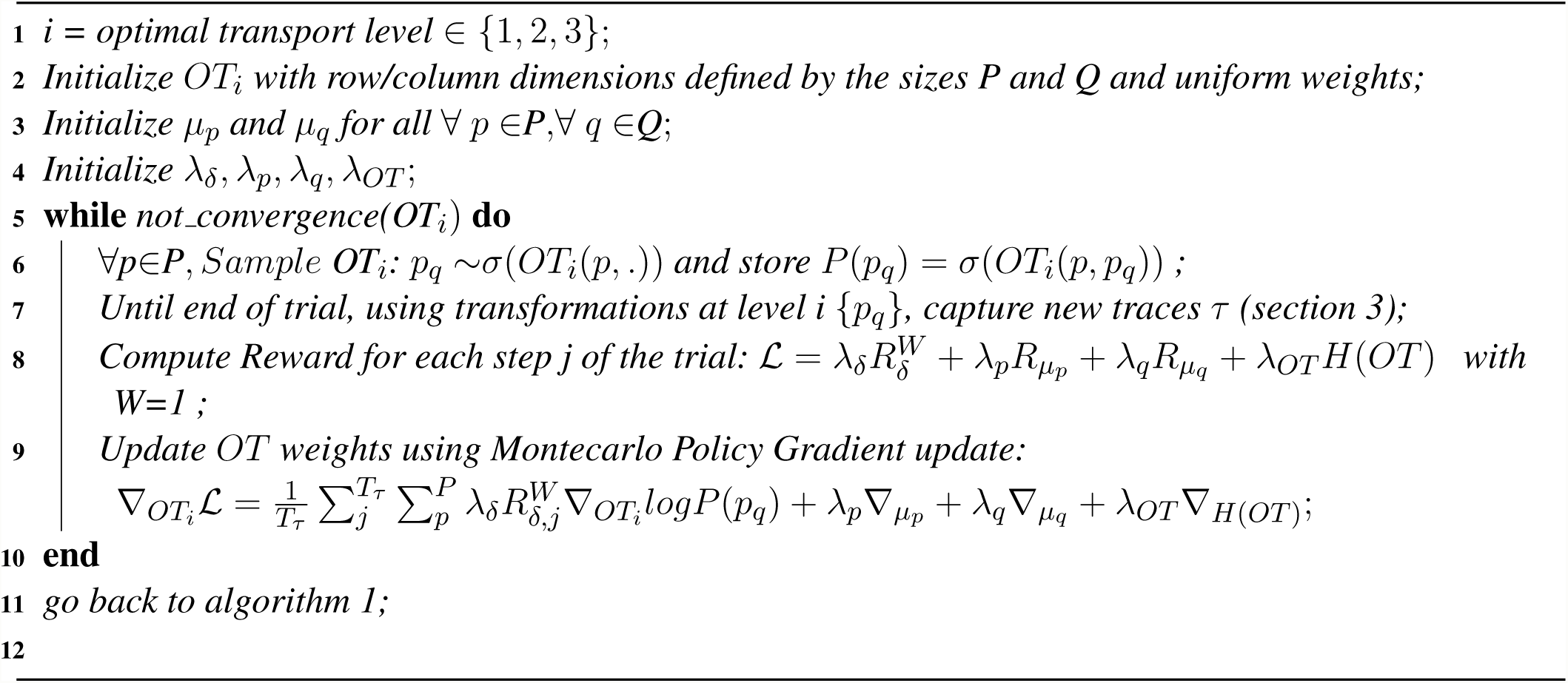

Algorithm 2 trains the weights of a transport matrix *OT*_*i*_ for a number of trials until convergence. Each trial, the algorithm generates a transport sample for each prototype vector of the new distribution (line 6): for each row of the *OT*_*i*_, assuming a categorical distribution with probability mass defined by *s*(*OT*_*i*_(*p*, .)), where *σ* is a softmax function, it generates a transport candidate *p*_*q*_. Line 6 finishes by storing the candidate probability (line 6). With a transport candidate for each prototype vector of the new input distribution, the agent will interact with the environment as discussed in section 3 (line 7). During the environment interaction phase, all *OT* functions will act as identity functions except the transport matrix under optimization *OT*_*i*_. For this case, each input *s*_*i,t*_ is first transformed to one of the *P* prototype vectors of the new input distribution and then transported to the original distribution using the current transport candidate *p*_*q*_. In line 8 the algorithm computes the components of the loss function defined in equation 6. Finally, line 9 computes the gradient updates. The gradients for 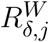 are computed using the REINFORCE update (Sutton and Barto, 2018) because this reward depend of a particular set of transport candidates sampled in line 6. Once the adaptation process (algorithm 1) finishes, the last optimized *OT*_*i*_ remains until algorithm 1 is activated again. Every activation of algorithm 1 starts with all *OT* functions acting again as identity functions.

### 3.4 Model experiments

The goal of this section is to validate the plausibility of the model. A canonical model of transfer learning does not exist (Haskell, 2000b), and as such we compare against a baseline model. On defining the baseline model, we specified the following modelling constraints: i) it should respect the multi-tiered architecture discussed in section 3, ii) the adaptation process should involve the least number of resources and iii) adaptation to a new task variant should be possible after learning a single task. Related to the second assumption, we would like to note that any change detection task variant with a maximum of 1 gray flash between images could be solved by uniquely optimizing the *DQN* tier because every unique image sequence of length three would probably be encoded in a unique way in the input of the *DQN* tier. In that sense, the simplest intervention (besides our optimal transport idea) to adapt to a new task variant would consist of uniquely optimizing the *DQN* tier. We did not consider weight fine-tuning strategies because the new *DQN* function could be drastically different than the original one (e.g. different image dictionary) and fine-tuning usually assumes pre-training with large number of task variants Cobbe et al. (2018) while we are assuming that our model has been pre-trained with a single task. Given the above discussion, our baseline model is based on a multi-tiered architecture pre-trained with training phase 1 and its adaptation process to new training phases consists of training the *DQN* tier starting from random weights ^5^.

Similarly to how we analyzed the transfer learning capacity in mice in section 2.1, we measure the learning efficiency when our model and the baseline model transition to training phase 2, 3 and 4 starting from a pre-trained state to solve training phase 1, 2 and 3 respectively. In all scenarios, our agent has been trained following the inference process described in section 3.2 with an environment that emulates training phase 1 or 2 or 3 as described in section 2 with a discrete time step corresponding to the 250 ms of a stimulus presentation and the 500 ms of a grey image presentation. For each case, we compute the time series of running *d*′^6^ four times and identify the first trial number with *d*′ greater or equal to 2. Finally, we quantify the models transfer learning ability by computing the rate defined in equation 2. In this case, besides the larger *d*′ threshold, rates are computed between every other pair of simulation runs because it does not exist the concept of mouse identifier. In every simulation we assume we know the correct tier to perform the representation remapping, so we are not using algorithm 1 because this is not a critical component of our quick adaptation proposal.

Table 3 shows the median and corresponding median absolute deviation of each 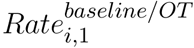 indicator. Row 1 summarizes the results for the baseline model and row 2 shows the results for our new model. We can observe that average baseline model rates are 3 to 4 times higher than average mice rates while the new model rates are of similar magnitude. Our model rates are not significantly bigger than mice rates with p-values^7^ of 0.105, 0.44 and 0.51 respectively, while baseline model rates are significantly bigger than mice rates with p-values^7^ of 8.62e-07, 0.000006 and 0.000014 respectively. Therefore, our model features a superior plausibility than the baseline.

**Table 3.** Median and median absolute deviation of 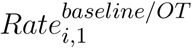 for i ∈ {2, 3, 4}. Each rate is computed for two cases (rows): a) baseline method based on learning uniquely the final tier *DQN* starting from random weights, b) proposed model based on distributed optimal transport. Statistics are computed from 4 simulations per case and rate indicator (We pick models which train well).

| Median (median absolute deviation) | $Ratio_{2,1}$ | $Ratio_{3,1}$ | $Ratio_{4,1}$ |
| --- | --- | --- | --- |
| Baseline | 1.27 ($\hat{\sigma} = 0.58$) | 0.96 ($\hat{\sigma} = 0.37$) | 1.65 ($\hat{\sigma} = 0.54$) |
| Optimal transport | 0.13 ($\hat{\sigma} = 0.05$) | 0.23 ($\hat{\sigma} = 0.08$) | 0.17 ($\hat{\sigma} = 0.04$) |

Figure 6 shows the model simulation results. Each plot corresponds to the evolution of *d*′ for 1000 trials averaged over 4 simulations starting with different random seeds. Blue curves show the agent adaptation when we train the final policy network *DQN*, while orange curves represent the agent adaptation when using the optimal transport mechanism before *Enc*1 (figures 6.b) and c)) or before *Enc*2 (figure 6.a)). Figure 6.a) corresponds to the case of changing the environment dynamics, Figure 6.b) corresponds to the case of changing the image dictionary to a new one with larger number of images (from 3 to 6), and finally figures 6.c) corresponds to the case of changing the image dictionary to a new one with the same number of images (3).

**Figure 6.**
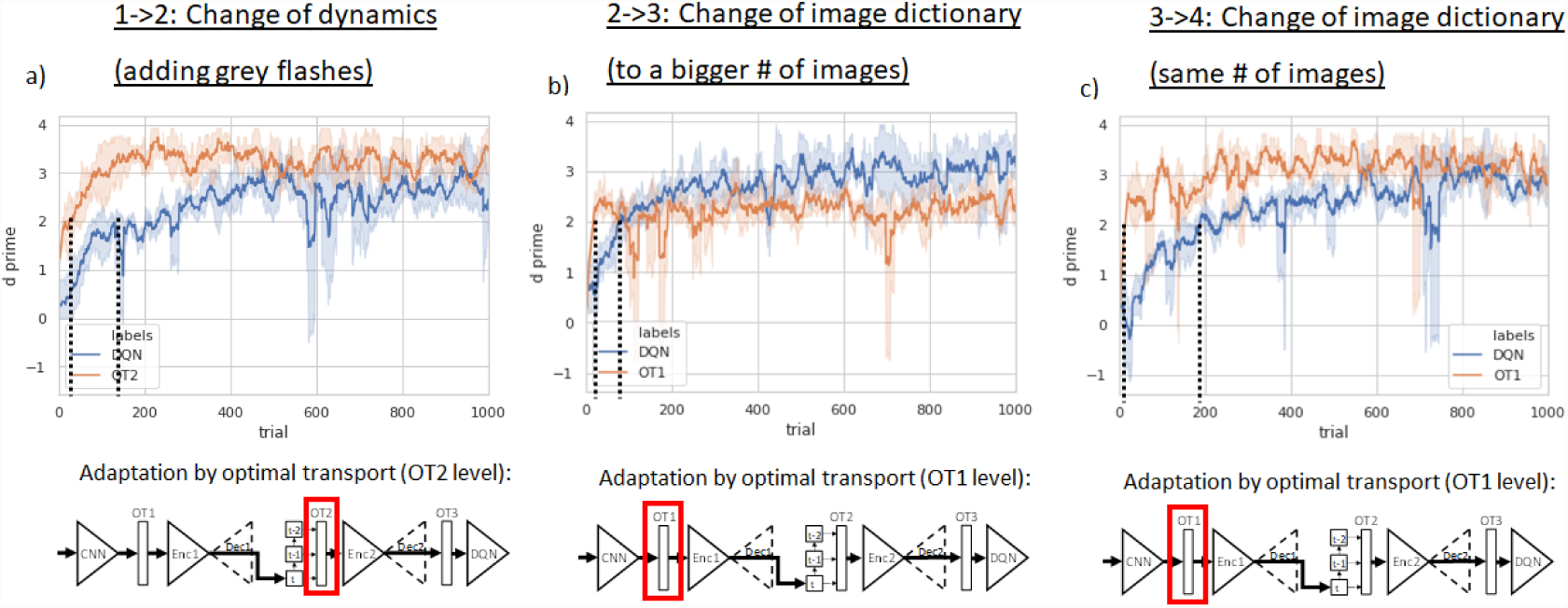
Model simulations.

Figure 6.a) shows that when there is a change in environment dynamics (e.g. adding grey flashes between image pairs), then this can be solved by performing an adaptation in *OT* 2. In this case, the agent is able to reach a *d*′ of 2 with an average of 15 trials 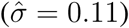 compared to the 163 trials 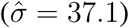 required by the reference case.

Figure 6.b) shows that our agent is also able to quickly reach a *d*′ of 2 with an average of 18 trials 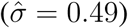 compared to the 81 trials 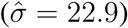 required when retraining the final policy network *DQN*. In this second case, algorithm 1 also chooses to optimize *OT* 1. However, the superior representation complexity required in the new image set limits the maximum *d*′ because few images from the new and larger image dictionary need to be mapped to the same images in the old dictionary. We hypothesize that the similar phenomena of reduced maximum performance observed in mice when moving to a task with larger representation complexity requirements can be due to similar cause. Precisely, mice may be capable of a quick adaption by recycling existing resources dedicated to similar tasks. We believe this mechanism is meant to increase survival probability when the environment conditions suddenly change by providing a temporal solution until a long term adaptation process is able to optimize the mouse behavior in the new environment.

Finally, figure 6.c) shows that our agent is able to reach a *d*′ of 2 with an average of 14 trials 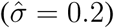 compared to the 187 trials 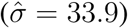 required when retraining the final policy network *DQN*. In this case, algorithm 1 is able to recover the agent operation by optimizing *OT* 1.

Implementation wise, *Enc*1, *Dec*1, *Enc*2 and *Dec*2 are implemented as feed forward neural networks with 2 hidden layers of 32 hidden units each and Relu activation units. Input dimensions of *Enc*1 and *Enc*2 are 84 and 18 respectively and output dimensions of *Enc*1 and *Enc*2 are 6 and 2 respectively. Policy network *DQN* is also a feed forward neural network with 2 hidden layers of 16 hidden units and Relu activation units. Input and output dimensions of *DQN* network are 6 and 2 respectively. All networks and the optimal transport matrix are trained using an Adam optimizer and a learning rate of 5*e*^−3^. The mini-batch size to train the policy network *DQN* and the auto-encoders was seto to 100, the *DQN* replay buffer size was 1000 and the probability of random action was constant and equal to 0.05. All networks were implemented with Pytorch.

Concerning the hyper-parameters of the lost function during the adaptation phase, we set them manually according to the following criteria. For the two use cases involving the optimization at the *OT* 1 level (figures 6.b and c), we used *λ*_*δ*_ = 0, *λ*_*p*_ = 0.05, *λ*_*q*_ = 0.05 and *λ*_*OT*_ = 2.0. We did not use 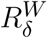 (equation 7) because in these two cases any mapping (*OT* 1) which preserves the probability masses of the old and the new stimulus distributions and minimizes entropy of *OT* 1 is valid. We believe that a similar heuristic may also be used by the mouse in order to minimize costly environment interactions. For the case involving the optimization at the *OT* 2 level (figure 6.a), we used *λ*_*δ*_ = 1.0, *λ*_*p*_ = 0.05, *λ*_*q*_ = 0.05 and *λ*_*OT*_ = 2.0. In this case, the validity constraints used in the previous two cases are also necessary conditions but not sufficient because not all valid mappings from an optimal transport point of view can recover the correct operation.

## 4 DISCUSSION

We showed that the visual behavior dataset from the Allen Brain Observatory is a feasible platform to study transfer learning in mice. We observe a significant increase of learning efficiency during the first session after transitioning to a new training phase compared to the first session of the first training phase. We also observe that transitioning to a similar task with superior representation complexity achieve a maximum average *d*′ slightly inferior than the maximum average *d*′ achieved in the last session of previous training phase. We also observe that transitions to more complex versions of the change detection task result in a performance decrease ^8^.

We have proposed a computational model based on two architectural principles: i) a multi-tiered architecture suggested by the well-known hierarchical organization of the sensory cortex and evidence from in-vivo recordings from monkeys trained on a change detection task; ii) the idea of distributed representation firewalls in the input of each tier responsible for the conversion of actual inputs to expected ones, thus preventing downstream encoding anomalies.

The model design was initially steered by functional requirements and just slightly constrained by neuroscience literature. This methodological approach guided us towards an implementation of the representation remapping based on optimal transport between probability distributions. Our implementation turned out to predict the reduced performance when mice move to a new more complex task due to limited representation resources which were optimized for the previous task. Similarly, the multi-tiered architecture was first an engineering constraint and later a biologic insight confirmed by the literature. A final utility factor of our design methodology came from the computations required to perform the representation remapping because those can be interesting indicators in future experiments when looking for neural correlates to confirm or refute our model.

We simulate our model to understand the adaptation capabilities to new task variants and we compare these results with respect to mice and with respect to a baseline model designed following also a multi-tiered architecture and a more traditional adaptation approach. This comparative study of the transfer learning ability in mice and machines is based on the computation of the relative learning efficiency when a mouse/model is exposed for the first time to training phases 2, 3 and 4 compared to the learning efficiency demonstrated during the first session of the first training phase. During the different types of task transitions we can observe that the average adaptation efficiency rates of the baseline model are 3 to 4 times higher than the average rates obtained in mice while the new model rates are of similar magnitude.

Finally, supplementary section 1 discusses a future research proposal. It extends our cognitive model and it is meant to contribute to both cognitive sciences and lifelong learning in artificial intelligence.

## Supporting information

Suplemental sections

## AUTHOR CONTRIBUTIONS

IM performed the analysis, implemented the model, performed the model simulations and wrote the paper. SM and IM designed the analysis and modelling studies. MG, DO, PG and SO contributed to design the experimental setup and to collect the data. SM, MG, DO, PG, SO and IM revised the manuscript.

## ACKNOWLEDGMENTS

We wish to thank the Allen Institute for Brain Science founder, Paul G. Allen, for his vision, encouragement and support.

## Footnotes

1 It spans 0.115 to 0.715 seconds, after accounting for a 35 ms monitor delay.

2 When the number of go trials is less than W, then we use this smaller number of go trials to compute the hit rate. When the number of catch trials is less than W, then we use this smaller number of catch trials to compute the false alarm rate.

3 Wilcoxon signed-rank test has the null hypothesis (one-sided) that the median of mice rates-1 is positive against the alternative hypothesis that it it is negative

4 During model inference we pick a random action with constant probability of *∈* = 0.05.

5 When we tried to re-train the *DQN* tier, starting from already trained weights, the agent was not able to recover its normal operation. We speculate that by training the agent on a single task, the *DQN* tier become to specialized to be fine-tuned to perform in the new task variant.

6 We compute *d′* according to equation 1 but with a smaller window *W* = 20 because the simulated agent learns and adapts faster than the mice.

7 Mann–Whitney U test (also called Wilcoxon rank-sum test) is a non-paired significance test. Null hypothesis (one-sided) is that median from mice rates is less or equal than the median model rates.

8 The maximum average *d′* obtained in the first session of the new training phase is slightly inferior than the maximum average *d′* achieved in the last session of previous training phase.

