## Supplementary material for "Quantifying and Modelling Transfer Learning in Mice Between Consecutive Training Stages of a Change Detection Task": Suplemental sections

### 1 MODEL GENERALIZATION

In this appendix we draft possible future model extensions to explain the quick adaptation phenomena in a large set of diverse environments. Our model generalization proposal is based on the idea that mice and other animals are capable of learning a grammar of the environment. In general, this grammar can be decomposed in a set of functional rules organized as a directed acyclic graph (DAG), where each rule input is a model input and/or any output from any other rule. Rule outputs feed other rules or it serve as quality value vector $q_t$. This environment grammar facilitate to the agent specialized sensory stream encoding, control and quick adaptation capabilities to succeed in a particular set of similar environments. This generalized architecture facilitates the internal knowledge structure required by an animal to learn and to transfer knowledge across similar tasks.

Figure S1 outlines the conceptual architecture of the generalized model. Just like the model discussed in section 3, each functional rule has three functionalities: i) feature extraction, ii) anomaly detection and iii) input domain adaptation through optimal transport. The first functionality or feature extraction does not change. The main change is in the anomaly detection. It is motivated by the requirement of being able to deal with environments where the agent actions affect future agent observations. Note that this was not the case in the change detection training setup discussed in section 2. Due to this new requirement, input distribution anomalies can arise due to changes in environment and/or changes in how the agent acts. Both cases can be captured using the prediction error of a forward model $\hat{s}'_{3,t+1} = f_3(s'_{3,t}, a_t)$ where $s'_{3,t}$ is the output of the optimal transport of rule 3 and $a_t$ is the agent action. Note that for some functional rules, the forward model may not depend of agent actions. The forward model gives also to the agent possibility to generate counter-factual encodings by testing the effects of alternative agent actions.

Figure S1 also shows that this model could be used to explain some illusions. For instance, it could explain why we seem to hear the electric tower jumps by simply watching the famous video without sound of an electric tower jumping. The optimal transport in node 3 would be responsible for transporting the images without sound to the corresponding images with sound that would reduce anomalies in node 3 and 4. This is indeed an alternative view of the predictive coding framework where discrepancies between bottom-up signal and top-down predictions drive anomalies which in turn are responsible for the selection of the node where an optimal transport intervention should take place. It suggests that prediction errors are not directly forwarded top-down but instead are used uniquely to drive adaptation.

Finally, we have been assuming during all this paper that the the environment grammar defined by the DAG exist but we didn't explained how. This is indeed an important research topic in itself which could shed light to how animals are able to accommodate to new tasks at adult age even though the brain plasticity is very limited during adulthood. We believe this could be due to meta-learning process responsible for reorganizing specialized resources which we encapsulate in the idea of functional rules. We plan to address this question in the near future.

### 2 TIME TO REACH D'=2 USING DIFFERENT REWARD FUNCTIONS

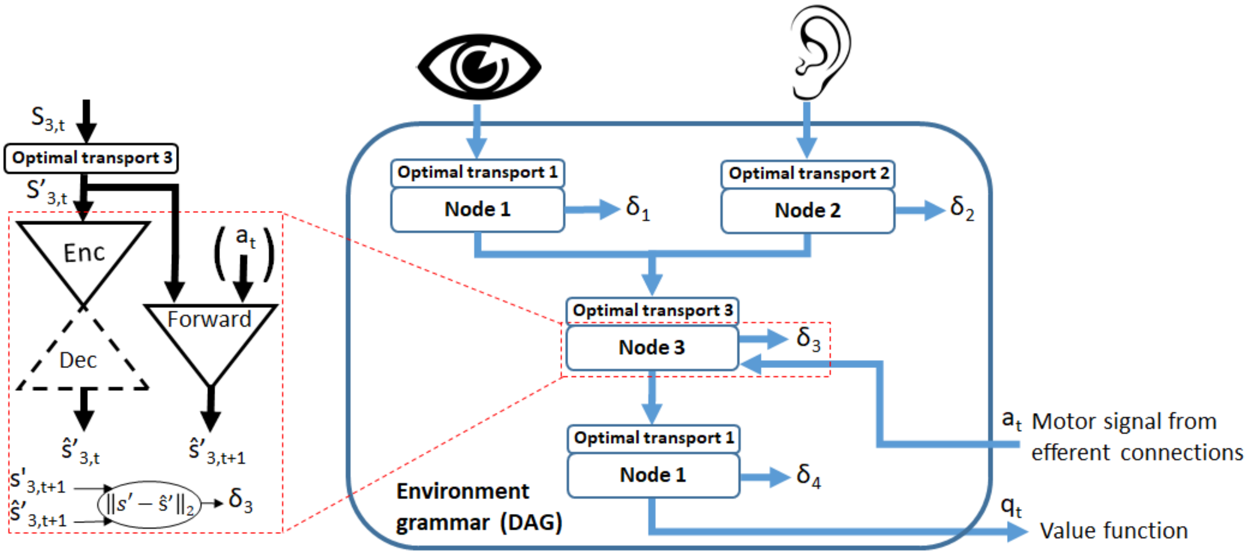


**Figure S1.** Conceptual architecture of the generalized model.

**Table S1.** Mean and standard deviation of the time required to reach d'=2 by training the DQN tier starting from random weights and using a dictionary of 3 images and 4 different reward functions (equation 11) which uniquely differ on the negative reward.

| $r_t$(Agent licks but not in last step of Go trial) $= -1.0$ | -0.2 | -0.1 | 0.0 |
|---|---|---|---|
| 229.0 ($\hat{\sigma} = 18.8$) | 228.0 ($\hat{\sigma} = 54.8$) | 179.7 ($\hat{\sigma} = 49.9$) | 325.1 ($\hat{\sigma} = 107.3$) |
